# Culmorin inhibits detoxification of the mycotoxin deoxynivalenol by plant UDP-glucosyltransferases

**DOI:** 10.1101/2025.11.06.686973

**Authors:** Herbert Michlmayr, Gerlinde Wiesenberger, Katrin Rehak, Marta Sopel, Kristina Funtak, Alexandra Malachová, Philipp Fruhmann, Julia Weber, Rudolf Krska, Marie Dufresne, Franz Berthiller, Gerhard Adam

**Affiliations:** BOKU University, Institute of Microbial Genetics, Department of Agricultural Sciences, Konrad-Lorenz-Strasse 24, 3430 Tulln, Austria; BOKU University, Institute of Crop Breeding and Genomics, Department of Agricultural Sciences, Konrad-Lorenz-Strasse 20, 3430 Tulln, Austria; BOKU University, Institute of Bioanalytics and Agro-Metabolomics, Department of Agricultural Sciences, Konrad-Lorenz-Strasse 20, 3430 Tulln, Austria; FFoQSI GmbH Austrian Competence Centre for Feed and Food Quality, Safety and Innovation, Technopark 1D, 3430 Tulln, Austria; Institute of Applied Synthetic Chemistry, TU Wien, Getreidemarkt 9, 1060 Vienna, Austria; Institute for Global Food Security, School of Biological Sciences, Queen’s University Belfast, 19 Chlorine Gardens Belfast BT9 5DL, Northern Ireland; Université Paris-Saclay, INRAE, UR BIOGER, 91120 Palaiseau, France

**Keywords:** Fusarium, glucosylation, inhibition kinetics, plant-pathogen interaction, root, secondary metabolism, sesquiterpene, trichothecene, virulence

## Abstract

The *Fusarium* metabolite culmorin (CUL) frequently co-occurs with the mycotoxin deoxynivalenol (DON) on cereals. While DON is recognized as a major *Fusarium* virulence factor on plants, the function of CUL is still unclear. Herein, we show that CUL-deficient *F. graminearum* mutants created by *CLM1* deletion are less aggressive on wheat than the wild-type, accompanied by increased DON-3-glucoside/DON ratios in infected wheat ears. In root elongation assays with wheat and *Brachypodium distachyon*, CUL had no effect alone but significantly increased the toxicity of DON. Analysis of DON/CUL-treated roots further indicated that both wheat and *B. distachyon* are able to glucosylate CUL and that its presence impedes DON-glucosylation in both species. We identified two *B. distachyon* UDP-glucosyltransferases (UGT) able to glucosylate CUL and further investigated the effect of CUL on the kinetics of validated DON-glucosylating plant UGTs (BdUGT5g03300, HvUGT13248, OsUGT79). This suggested that CUL inhibits DON-glucosylation either by serving as competitive substrate with DON or by unproductive binding. Especially BdUGT5g03300 was strongly inhibited by CUL and even its glucosides. Our results indicate that CUL contributes to *Fusarium* virulence by weakening plant-defenses related to UGT-catalyzed DON-detoxification. As even CUL-glucosides are potentially inhibitory to UGTs, this implies a complex synergy of CUL with DON.

**Highlight:** We present biochemical evidence that the *Fusarium* metabolite culmorin contributes to *Fusarium* virulence on plants by suppressing detoxification of the virulence factor deoxynivalenol by glucosyltransferases.

## Introduction

Infection by *Fusarium* species is a global threat to cereal crops such as wheat, barley and maize, causing drastic yield-losses and quality deterioration due to mycotoxin contamination (Chen *et al*., 2019; Goswami and Kistler, 2004). The (sesquiterpene) trichothecene mycotoxin deoxynivalenol (DON) inhibits eukaryotic protein synthesis and is a crucial virulence factor of *Fusarium graminearum*, required for spreading in wheat ears (Bai *et al*., 2002; Desjardins *et al*., 1996; Proctor *et al*., 1995). DON is a highly problematic food contaminant and due to its ubiquitous occurrence one of the economically most important mycotoxins. Acute or chronic exposure cause several clinical symptoms in humans and animals (Pestka, 2007; Pestka, 2010). Legislations or guidance values have therefore been enacted for instance in the United States (U.S. Food & Drug Administration, 2010), the European Union (The European Commission, 2023) and several Asian countries (Anukul *et al*., 2013).

The tricyclic sesquiterpene diol culmorin (CUL) is produced by several *Fusarium* species including *F. graminearum, F. crookwellense, F. venenatum* and the name-giving *F. culmorum* (Pedersen and Miller, 1999). Upon its first description (Ashley *et al*., 1937), it received little attention due to its low apparent toxigenic properties. However, several studies indicated that CUL persistently co-occurs with DON in equivalent or even higher concentration levels (Ghebremeskel and Langseth, 2001; Hajnal *et al*., 2020; Radić *et al*., 2021; Streit *et al*., 2013; Uhlig *et al*., 2013), raising concerns towards its function and toxicological potential. Weak antifungal activity of CUL against several marine and terrestrial (human pathogenic) fungi has been reported (Strongman *et al*., 1987). *In vitro* assays further implied low toxicity to animal cells (Gruber-Dorninger *et al*., 2017), no negative effects were observed on the growth of piglets (Rotter *et al*., 1992) and *Heliothis zea* larvae (Dowd *et al*., 1989). However, CUL significantly increased mortality of the latter when applied in combination with DON. The authors were the first to propose a synergistic effect with DON due to competition for detoxification enzymes (Dowd *et al*., 1989).

As sesquiterpenes, trichothecene class toxins (such as DON) and CUL share farnesyl diphosphate as biosynthetic precursor. The trichodiene synthase encoded by *TRI5* catalyzes the first step in trichothecene biosynthesis by converting farnesyl diphosphate to trichodiene (McCormick *et al*., 2011). Disruption of *TRI5* in *F. graminearum* abolished DON synthesis and resulted in reduced virulence (Proctor *et al*., 1995). Synthesis of culmorin occurs in two steps, starting with the conversion of farnesyl diphosphate to longiborneol by the terpene synthase Clm1 (McCormick *et al*., 2010). The cytochrome P450 Clm2 catalyzes the subsequent hydroxylation of longiborneol to CUL (Bahadoor *et al*., 2015). Both *CLM1* and *CLM2* are induced under trichothecene inducing conditions (Harris *et al*., 2016; McCormick *et al*., 2010; Sieber *et al*., 2014; Wipfler *et al*., 2019). Although this suggests a role in virulence, there is scarce evidence for that. A phytotoxic effect of CUL was reported on wheat coleoptile tissue at high concentration levels (1000 µM CUL) (Wang and Miller, 1988). However, at concentrations closer to natural contaminations levels (10 µM ≈ 3 mg L^-1^ range) CUL had no phytotoxicity on wheat by itself but synergistically increased the toxicity of DON and several other trichothecene toxins (Wang and Miller, 1988). Wipfler *et al*. (2019) reported synergistic effects of CUL and DON on wheat, barley and corn. Fusarium head blight disease severity on wheat caused by 15 *F. graminearum* isolates was positively correlated with the sum of CUL and DON but negatively correlated with the CUL/DON ratio (Wipfler *et al*., 2019).

The aim of the present study was to investigate the effect of CUL on wheat and *Brachypodium distachyon*, in particular by addressing its potential interaction/synergy with DON. Therefore, we created CUL-deficient *F. graminearum* strains to test whether the (in)ability to synthesize CUL has an impact on virulence on wheat. We further investigated the combined effect of CUL and DON on wheat and *Brachypodium distachyon* roots. These experiments implied that CUL impairs detoxification of DON to DON-3-glucoside (D3G) in plants. We further conducted experiments to investigate whether CUL can biochemically interfere with the detoxification of DON by previously validated DON-detoxifying plant UDP-glucosyltransferases (UGTs) in steady state kinetic assays.

## Materials and Methods

### Materials

DON was purified as described by Altpeter and Posselt (1994). CUL and its conjugated derivatives were synthesized and purified as described in Weber *et al*. (2018).

### Fungal strains, cultivation and construction of knock-out strains

*Fusarium graminearum* PH-1 (NRRL 31084, FGSC 9075) was routinely grown on Fusarium Minimal Medium (Leslie and Summerell, 2006) and sporulated in mung bean broth (10 g mung beans per litre of water). Conidiospores were quantified by counting them in a Fuchs-Rosenthal chamber. For determination of mycotoxins, strains were cultivated in 2-SM medium (3 g L^-1^ KH_2_PO_4_, 0.2 g L^-1^ MgSO_4_^*^7 H_2_O, 5 g L^-1^ NaCl, 1 g L^-1^ (NH_4_)_2_HPO_4_, 40 g L^-1^ sucrose, 10 g L^-1^ glycerol, 10 mg L^-1^ citric acid, 10 mg L^-1^ ZnSO_4_^*^6 H_2_O, 2 mg L^-1^ Fe(NH_4_)_2_(SO_4_)_2_^*^6 H_2_O, 0.5 mg L^-1^ CuSO_4_ ^*^5 H_2_O, 0.1 mg L^-1^ MnSO_4_, 0.1 mg L^-1^ H_3_BO_4_, 0.1 mg L^-1^ Na_2_ MoO_4_^*^2 H_2_ O). Magenta™ vessels containing 20 mL of 2-SM medium were inoculated with 10^5^ spores and incubated at 20 °C in the dark for three weeks. Mycelia were separated from medium using a Buchner funnel fitted with a filter paper and flow-through was analyzed by HPLC-MS/MS.

For deletion of the *CLM1* (FGSG_10397) gene in PH-1 two constructs were produced. First, the 3’ flanking region of the gene was amplified with the primer pair AAT<u>GTCGAC</u>TATGACAAACCAGATCTGAGTAAC and TAC<u>AAGCTT</u>GAAGACGATCAGCTGCATCCTA, the resulting amplicon was digested with SalI and HindIII and the 532 bp fragment was the ligated to pKT245 (Twaruschek *et al*., 2018) cleaved with the same enzymes. The resulting plasmid was termed pKF33. Similarly, the 5’ flanking region was amplified using primers TTATCTATTA<u>GGCCGCATAGGCC</u>TGATTG and TAA<u>ACTAGT</u>GTCTATATTCCGGCTGTAGATCA and the fragment was cloned into pKT245 via SfiI and SpeI, yielding pKF65. Next, pKF33 was digested with HindIII, RsrII and SacII, and pKF65 with SfiI and PstI to release fragments of 3128 bp and 2074 bp, respectively. Fragments containing either the 3’- or the 5’-flanking region plus non-functional parts of the *nptII* selection marker were mixed in a 1:1 molar ratio and 10 µg of this DNA mix were used to transform PH-1 (as described in Twaruschek *et al*. (2018), selecting transformants on 40 mg L^-1^ Geneticin (G418). Five transformants with the correct DNA integration (clm1Δ::*loxP*-HSVtk-*nptII-loxP*) were identified by PCR analysis, two of these were used for further work (KFCUL-37 and KFCUL-41). By treating these strains with CRE-recombinase the nptII marker was removed yielding strains GWCUL-37.37 and GWCUL-41.41 (*clm1*Δ::*loxP*). PH1t was regenerated from diluted protoplasts of a mock transformation (no DNA, no selective agent).

### Wheat infection

For wheat infection experiments Apogee wheat was grown in soil (20 °C, 22 hours light, 2 hours dark) until flowering. Fusarium infection was carried out by inoculating four flowers/ear with 10 µL each of a spore suspension containing 4^*^10^4^ conidia mL^-1^. 10 ears were treated per strain (PH-1, PH-1t, GWCUL-37.37 and GWCUL-41.41). The infection experiment was performed three times independently. Starting at day four post infection, progression of infection symptoms was assessed every other day until day 10 or day 12, respectively. On the last day, the wheat ears were cut off the stems and frozen in liquid nitrogen and stored at -80 °C until work-up.

Wheat ears were ground using a Retsch Ball Mill MM 400 (Retsch GmbH, Hann, Germany) and aliquots of the powder were weighed into 1.5 mL tubes. A 10-fold volume of 50% MeOH was added and samples were extracted on a Vibrax shaker (IKA, Staufen, Germany) for 30 minutes at room temperature. After centrifugation (15 minutes, 21,130 g, 4 °C) samples were appropriately diluted with methanol/water (1:1) and analyzed by HPLC-MS/MS.

### Root elongation assays

Murashige-Skoog (MS) (Murashige and Skoog, 1962) was prepared without sucrose. Apogee wheat (Bugbee *et al*., 1997) seeds were sterilized with 1.4% (w/v) NaOCl, and 0.06% (*w/v*) Triton-X-100 for 10 min under shaking. Afterwards, the seeds were washed twice with sterile H_2_O. For germination, the seeds were placed on wet (9 mL 1/2 x MS without sucrose) filter papers in 145 x 20 mm petri dishes and incubated for 24 h in a growth chamber set to 20 °C, 75 % relative humidity and a cycle of 18 h light and 6 h dark.

Growth assays were performed in 1/2 x MS without sucrose with 0.5% (w/v) agarose (SERVA Electrophoresis GmbH, Heidelberg, Germany) containing DON and/or CUL at 5 mg L^-1^. CUL was dissolved in DMSO, therefore, all media including the controls were adjusted to the same DMSO concentration of 1% (v/v). The assays were done in 200 µL pipette tips (200 µL wide orifice, DENVILLE Scientific, Metuchen, NJ) filled with 360 µL agarose medium. The tips were placed into flat bottom micro-inserts (0.2 mL, 31 x 6 mm) for HPLC vials (VWR, Darmstadt, Germany). The Apogee seedlings were “planted” into the agarose when the roots were clearly developed but less than 5 mm long, then grown at 20 °C, 75% relative humidity with a light-dark cycle of 18 h light and 6 h dark. Photographs of the experimental set-up are shown in **Supplementary Fig. S1**. After 96 h, the roots were cut from the plants, washed to remove agarose residues and total root length was analyzed with the WinRHIZO Pro 2013 software (Regent Instruments Inc., Québec, Canada) at a scanning resolution of 400 dpi (EPSON PERFECTION V700 PHOTO).

*B. distachyon* seeds were sterilized with 0.7% sodium hypochlorite (NaOCl) for 20 min by gentle shaking (80 rpm) at room temperature, then washed three times with sterile H_2_O. The seeds were then stratified in the dark at 4 °C for 1-2 weeks. For germination, the seeds were placed on (sterile) damp filter in 145x20 mm petri dishes, kept further at 4 °C (dark) for three days, and then placed at 20 °C in the dark. *B. distachyon* seedlings were transferred to agarose when the primary roots were about 5 mm long. The agarose concentration was reduced to 0.3% to make it easier to penetrate the medium without injuring the roots. Most plants developed one to three, but in the most cases, two adventitious roots. Total root length was analyzed with WinRHIZO as specified above.

### Root extracts

Apogee and *B. distachyon* seedlings with 1-2 cm long roots were incubated in 1 x MS without sucrose containing 50 mg L^-1^ DON and/or 50 mg L^-1^ CUL. The seedlings were placed into sterile 2 mL tubes containing 500 µL medium such that only the roots were submerged. After 24 h incubation under conditions as specified above, the roots were cut from the seedlings, washed with 10% ethanol, shortly dried on tissue paper and weighed. Afterwards the roots were ground in the Retsch Ball Mill MM 400 for 1 min at 30 x s^-1^ using two 5 mm steel balls. 500 µl methanol was added and the roots were extracted for 2 h at room temperature. The concentrations of DON, D3G, CUL and CUL-glucoside were determined by LC-MS/MS (see below).

### LC-MS/MS determination of DON, CUL and their glucosides

DON, D3G, CUL and CUL-glucosides were measured on a 6500+ QTrap MS/MS instrument (Sciex, Framingham, USA) coupled to a 1290 UHPLC system (Agilent, Waldbronn, Germany). A Kinetex C18 column (150 x 2.1 mm, 2.6 µm particle size) was used for separation at 30 °C (Phenomenex, Torrance, USA). Mobile phases consisted of mixtures of methanol and water (A: 10% MeOH, B: 97% MeOH) and both contained 5 mM ammonium acetate. A flow rate of 250 µL/min and an injection volume of 1 µL were chosen. The gradient started with 0% B for 1.0 min and linearly to 100% B at 8.5 min. The column was rinsed with 100% B for 3 min, before equilibration with the start conditions for another 3 min till the end of the run at 14.5 min. The switching valve of the mass spectrometer directed the eluent into the ion source between 3.0 and 11.0 min. The IonDrive Turbo V electrospray source (Sciex) was operated at 550 °C with 50 psi of gas 1 and gas 2, 30 psi curtain gas and ionization voltages of +4500 V in both positive and -4500 V negative mode. DON and D3G were detected in negative ion mode first, while after 4.5 min polarity was changed to positive electrospray mode to detect CUL and CUL-glucosides The dwell time for each selected reaction monitoring (SRM) transition was set to 50 ms with a pause time between transitions of 5 ms. The following SRM transitions were used for CUL: m/z 256.2 > 221.2 (collision energy (CE) 17 eV) and m/z 256.2 > 203.2 (CE 21 eV) using the [M+NH_4_]^+^ ion as precursor with a declustering potential (DP) of 70 V. For CUL-glucosides also the [M+NH_4_]^+^ ion was used as precursor (DP 70 V) with the following SRM settings: *m/z* 418.2 > 177.3 (CE 35 eV) and *m/z* 418.2 > 221.2 (CE 30 eV). For DON the [M+CH_3_COO]^-^ ion was used as precursor (DP -70 V) with the following SRM transitions: *m/z* 355.1 > 265.1 (CE -22 eV) and m/z 355.1 > 59.1 (CE -40 eV). D3G was determined with its [M+CH_3_COO]^-^ ion as precursor (DP -80 V) with the following SRM settings: *m/z* 517.2 > 427.1 (CE -30 eV) and m/z 517.2 > 59.1 (CE -85 eV). Retention times were 3.21 min (D3G), 3.37 min (DON), 8.31 min (CUL-glucosides) and 8.91 min (CUL). Using these chromatographic conditions, CUL-8-glucoside and CUL-11-glucoside could not be separated.

### Glucosyltransferase assays

OsUGT79 (NM_001058779), BdUGT5g03300 (KQJ81826.1) and HvUGT13248 (ADC92550.1) were expressed with *E. coli* and purified as described (Michlmayr *et al*., 2018). Assays with DON and CUL were conducted at 25 µM substrate concentration at 25 °C and contained 10 mM UDP-glucose (Uridine 5⍰-diphosphoglucose disodium salt hydrate from *Saccharomyces cerevisiae*; Sigma-Aldrich, Vienna, Austria) and 100 mM potassium phosphate buffer pH 7.0. After 10-15 min, the assays were stopped by adding a 10-fold volumetric excess of methanol. Kinetics of HvUGT13248 with CUL were assayed under above mentioned conditions using 5, 12.5, 25, 50, 125 and 250 µM CUL. The stopped samples were stored at 4 °C until DON, D3G, CUL and CUL-glucoside were quantified by LC-MS/MS after 1:10 dilution in methanol. Reaction rates were calculated as µmol or nmol glucoside formed per min and mg of protein. All assays were conducted in triplicate.

### Steady state kinetics and kinetic models

Steady state enzyme kinetics were measured with a coupled enzymatic reaction to detect released UDP *via* NADH oxidation (Wetterhorn *et al*., 2017). The reaction contained 100 mM Tris·HCl (pH 7.0), 2 mM phosphoenolpyruvate (Roche, Basel, Switzerland), 0.2 mM NADH (Roche), 5 mM MgCl_2_, 2 µL (in total assay volume of 40 µl) pyruvate kinase/lactic acid dehydrogenase from rabbit muscle (Sigma-Aldrich) and 1 mM UDP-glucose. If necessary, UGTs were diluted in 100 mM Tris·HCl (pH 7.0) in order to obtain a linear time-dependent response. CUL was dissolved in methanol and added to the reaction mix from a 50-fold stock. Reactions were started by addition of DON dissolved in H_2_O, blanks contained H_2_O instead of DON. Assays were done in 384 well plates, and the reaction was monitored by reading the absorbance at 340 nm in an EnSpire 2300 Multilabel Reader (PerkinElmer, Waltham, MA, USA) adjusted to 25 °C. Enzyme reaction rates were calculated from NADH decrease using a NADH calibration curve measured under similar conditions (pH, buffer, temperature) and expressed as µmol min^-1^ mg^-1^ referring to release of UDP per mg of one-step purified protein.

The paramaters *V*_max_ and *K*_M_ were estimated by iterative curve fit with SigmaPlot 11.0 (Systat Software, San Jose, CA, USA) using the Michaelis-Menten model (equation 1) or the Haldane model of substrate inhibition (equation 2). Inhibition by culmorin was analyzed by a classic approach using Lineweaver-Burk diagrams following Bisswanger (2015). The corresponding models of competitive, uncompetitive and non-competitive (mixed) inhibitions are shown in equations 3, 4 and 5, respectively. The inhibitor constants *K*_ic_ (competitive) and *K*_iu_ (uncompetitive) were estimated from secondary diagrams plotting slopes and y-axis intercepts of the primary Lineweaver-Burk regressions versus inhibitor concentration [I]. The x-intercept (y=0) of the regression in the secondary plot corresponds to the respective negative value of the respective *K*_i_: Plotting the slopes versus [I] yields -*K*_ic_, plotting y-intercepts versus [I] yields -*K*_iu_.

1. Michaelis-Menten model 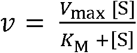
2. Substrate inhibition 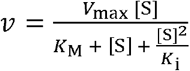
3. Competitive inhibition 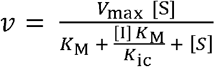
4. Uncompetitive inhibition 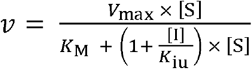
5. Full kinetic model of mixed inhibition 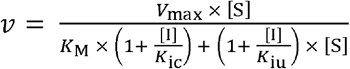

### Statistical analysis

All statistical tests were performed in R Studio. Infection scores and root length assays were tested for normality with the Shapiro-Wilk test, homogeneity of variances was tested with the Levene test implemented in the car package for R (Fox and Weisberg, 2019). Since each dataset contained at least one group that deviated from normal distribution (judged by Shapiro-Wilk p-value), all datasets were analyzed with the Kruskal-Wallis rank sum test implemented in R. Post hoc analysis for pairwise comparisons were conducted with the Wilcoxon rank sum test using the Benjamini – Hochberg method (Benjamini and Hochberg, 1995) to adjust for multiple testing.

Paired t-test (two-sample, two-tailed) was used to compare analytical results.

## Results

### Deletion of *CLM1* reduces the virulence of *F. graminearum* PH-1 on wheat

To investigate whether CUL plays a role in virulence, we constructed *F. graminearum* mutant strains (in the PH-1 background) by deleting the longiborneol synthase gene *CLM1* (FGSG_10397). CUL production was completely abolished in *clm1* deletion strains while these produced slightly higher amounts of DON than the wild type PH-1 on 2-SM medium (**Supplementary Table S1**). The subsequently generated marker-less *clm1*::*loxP* mutant strains GWCUL-37.37 and GWCUL-41.41 were used for further experiments. Apogee wheat was infected with these strains, the wild-type *Fusarium graminearum* (PH-1) and a transformation negative control (PH-1t) in three independent infection experiments (**Fig. 1**). All three experiments indicated that the CUL-deficient mutant strains are less aggressive on wheat than the parental PH-1. Pairwise statistical analyses showed that not all comparisons (mutant vs. wild type) were significantly different at the α = 0.05 level. Nevertheless, the overall results of the three independent replications clearly suggest that the *clm1* mutants differ from the PH-1 controls (**Supplementary Table S2**). At the end of two infection experiments (**Fig. 1 BC**), wheat ears were harvested, extracted and analyzed for their contents of DON, D3G, CUL and CUL-glucosides (the used analytical method could not distinguish between CUL-8- and CUL-11-glucoside). The ears infected with the *clm1* deletion strains did not contain any CUL or CUL-glucoside, while both compounds were detected in wheat ears infected with PH-1 (**Supplementary Table S3**). Evaluation of the DON and D3G contents in wheat ears revealed a higher DON content in the wild type infections, while D3G concentrations were comparable across all experiments. However, the D3G/DON ratios were significantly increased in *clm1*-infected wheat ears compared to those infected with the wild-type strain PH-1, indicating an interference of CUL with the glucosylation of DON in wheat (**Supplementary Table S3**).

**Fig. 1.**
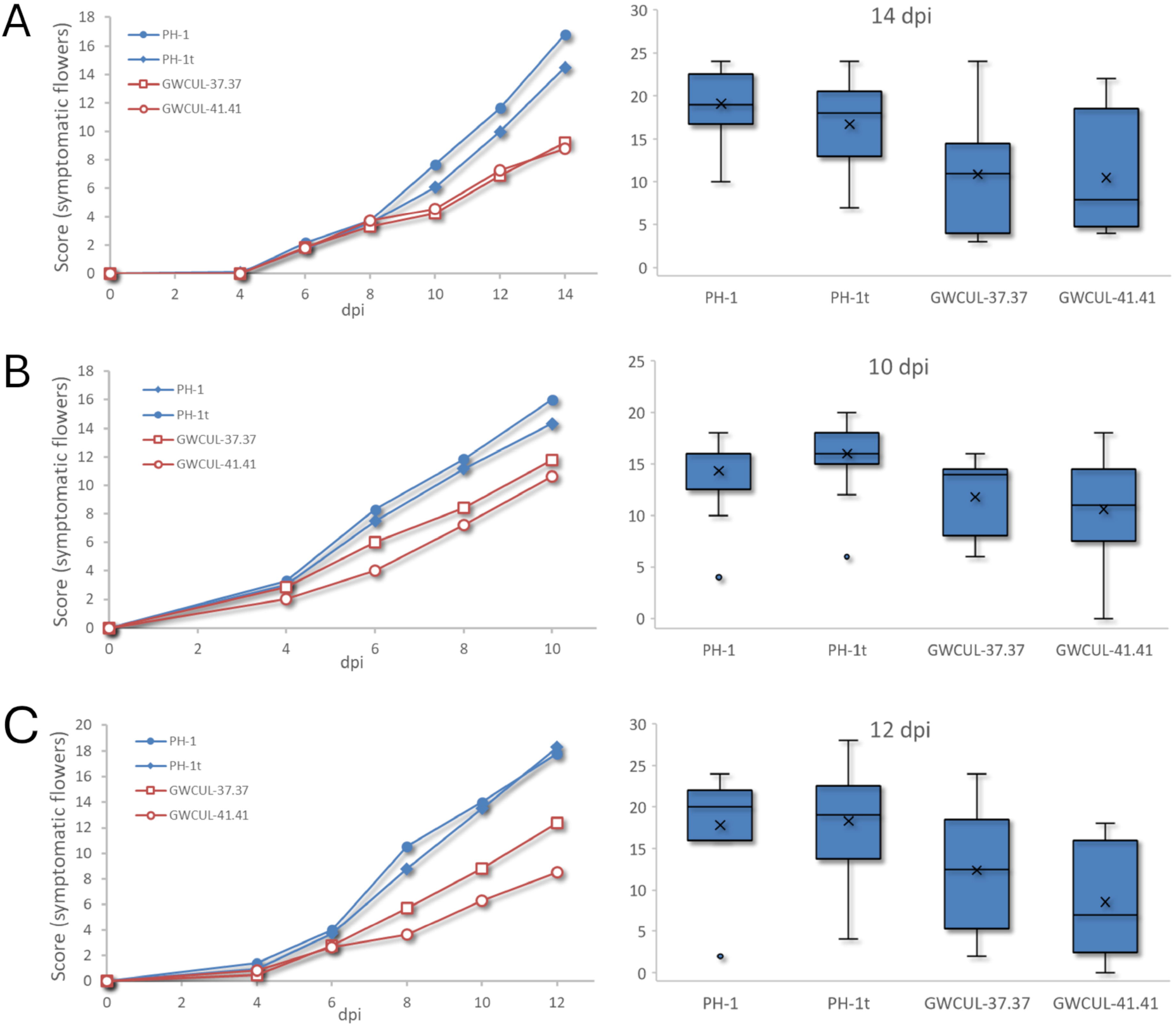
Disease progression of Apogee wheat ears infected with Fusarium graminearum wild-type strains (PH-1, PH-1t) and CUL deficient clm1Δ mutants GWCUL-37.37 and GWCUL-41.41. Shown are average scores (n = 10) of symptomatic flowers of three independent experiments (A, B, C). The left panel shows disease progression by score average (dpi, days post infection), the left panel shows boxplots of the respective last time-points, x indicates the score average. Statistical analysis of the last data points (Wilcoxon test) is shown in Supplementary Table S2.

### Culmorin interferes with DON-glucosylation in wheat roots

Root elongation assays were previously used as toxicity assays for DON and other trichothecenes in *A. thaliana* (Masuda *et al*., 2007), *B. distachyon* (Pasquet *et al*., 2016) and wheat (Wipfler *et al*., 2019). We compared the root length of Apogee wheat seedlings after 96 h of growth on agarose medium containing CUL and/or DON (**Fig. 2 A-C**). DON caused a significant reduction of both primary (longest root) and total root length (statistics in **Supplementary Tables S4&5**). While CUL alone did not cause a significant change, application together with DON increased the toxicity of DON significantly by causing a further reduction of root length (**Fig. 2, Supplementary Tables S4&5**), confirming previous observations (Wipfler *et al*., 2019).

**Fig. 2.**
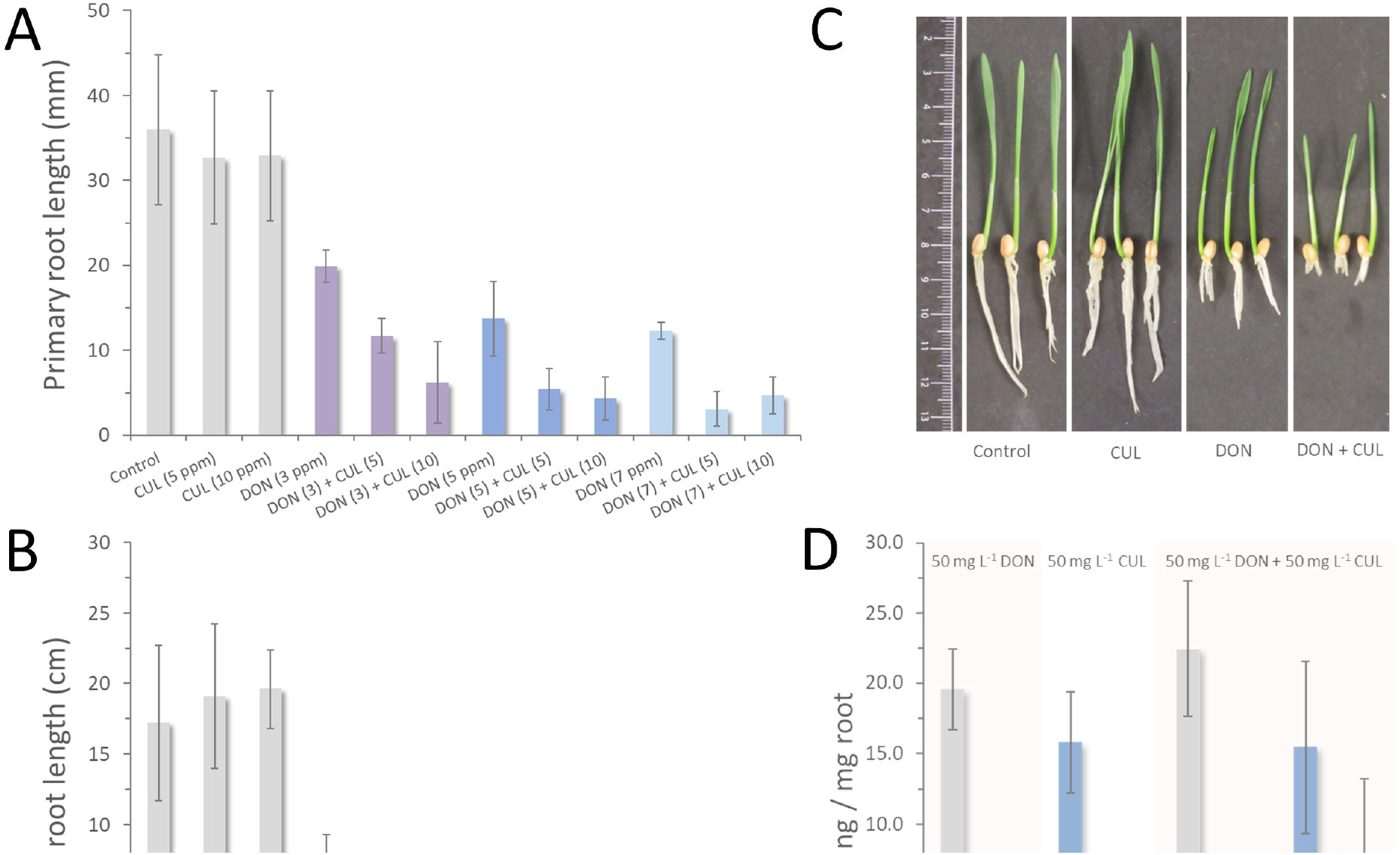
Impact of deoxynivalenol (DON) and culmorin (CUL) on roots of wheat (cv. Apogee) seedlings. **(A)** Average maximum root length (n = 10, error bars indicate SD) of Apogee seedlings after 96 h growth in Murashige-Skoog (MS) agarose medium containing different concentrations of DON and CUL. Concentrations in mg L^-1^ are given in parentheses. **(B)** Average total root length of Apogee seedlings under conditions as specified in **(A). (C)** Apogee seedlings photographed after 96 h growth in MS agarose medium containing 5 mg L^-1^ DON and/or CUL. **(D)** DON, DON-3-glucoside (D3G), CUL, CUL-glucoside (CULG) in Apogee roots extracted after 24 h incubation in MS with 50 mg L^-1^ (169 µM) DON and/or 50 mg L^-1^ (210 µM) CUL. The displayed results are means of three biological replicates ± standard deviation.

Analysis of Apogee root extracts prepared after incubation of seedlings with DON and/or CUL confirmed a previous report (Weber *et al*., 2018) that wheat is able to glucosylate CUL (**Fig. 2D**). The presence of CUL caused an approximately 6-fold reduction (n = 3; t-test, *p* = 0.05,) of D3G levels in the roots, suggesting that the synergy observed with DON is related to impaired ability to detoxify DON by UGTs.

### Culmorin inhibits DON-conjugation by UGTs

The above results indicate that CUL in fact contributes to virulence and that its presence has a negative impact on DON-glucosylation by wheat UGTs. A question is therefore whether CUL interferes with DON detoxification by acting as a potential competitive substrate or inhibitor of UGTs. In order to test whether CUL interacts with UGTs, we investigated the previously characterized UGTs HvUGT13248 (barley), BdUGT5g03300 (*B. distachyon*) and OsUGT79 (rice), known to efficiently detoxify DON by glucosylation at position C3 (Michlmayr *et al*., 2018).

Simultaneous incubation of DON with CUL or CUL-conjugates (equimolar or in five-fold excess over DON) clearly showed that CUL and its conjugates reduce DON-glucosylation rates of all the investigated UGTs (**Table 1**). The D3G synthesis rates of HvUGT13248 and OsUGT79 were reduced by CUL and CUL-11-acetate while the glucosides and sulfates had moderate to no effect. BdUGT5g03300 was almost inactive (rate reduced to < 1%) when CUL and CUL-11-acetate were present equimolar with DON. Even the glucosides and sulfates (except for CUL-8,11-disulfate) had a strong inhibitory effect on DON-glucosylation by BdUGT5g03300.

**Table 1.**
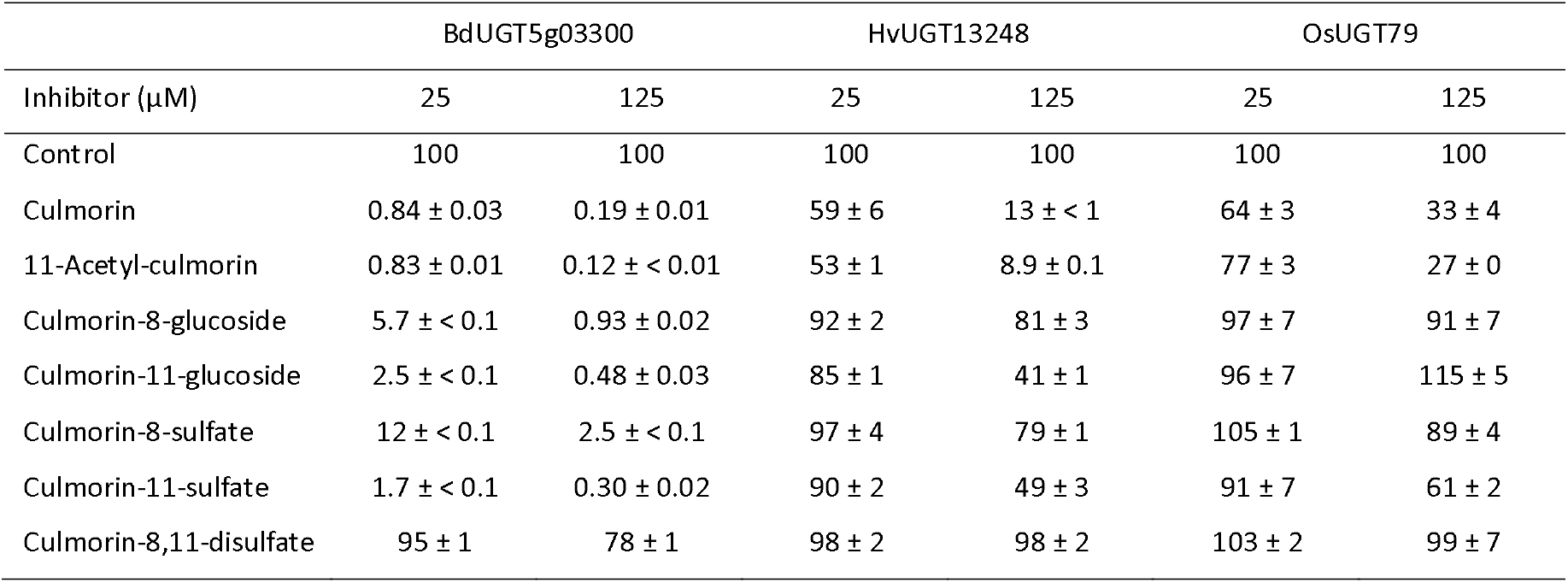
Glucosylation of deoxynivalenol (DON) by UDP-glucosyltransferases in presence of culmorin (CUL) and culmorin-conjugates. CUL and CUL-conjugates were applied equimolar (25 µM) or in five-fold molar excess (125 µM) over DON (25 µM in assay). Assays were performed at 25 °C, 10 mM UDP-glucose, pH 7. DON-3-glucoside synthesis was quantified by liquid chromatography/tandem mass spectrometry. The results are expressed as reaction rates (%) compared to the controls (100%) without inhibitor and represent the average of triplicate determination ± standard deviation.

### Culmorin glucosylation by UGTs

To clarify whether reduction of activity with DON is a result of CUL competing with DON as substrate, we compared the specific activities of these UGTs with DON and CUL at 25 µM each (**Table 2**). This indicated that BdUGT5g03300 is almost inactive with CUL. Even after several hours of incubation, only traces of CUL-glucoside were detectable by liquid chromatography coupled to tandem mass spectrometry (LC-MS/MS). Specific activity with CUL was (roughly) estimated thousand-fold below the value obtained with DON. By contrast, HvUGT13248 glucosylated both compounds at comparable rates. Further kinetic analysis of HvUGT13248 with CUL revealed high affinity for CUL with a *K*_M_ of 27 ± 4 µM CUL and a *V*_max_ of 18.2 ± 0.8 nmol min^-1^ mg^-1^ (**Supplementary Fig. S2**). Judged by comparing *V*_max_ /*K*_M_ ratios, catalytic efficiency of HvUGT13248 with CUL is comparable or even higher than with DON (*K*_M_ = 1.8 mM DON and *V*_max_ = 0.18 µmol min^-1^ mg^-1^).

**Table 2.** Specific activities of UDP-glucosyltransferases with deoxynivalenol (DON) and culmorin (CUL) at 25 µM, 25 °C, 10 mM UDP-glucose in 100 mM potassium phosphate buffer pH 7. The results refer to glucoside formation as quantified by LC/MS and are expressed as nmol min^-1^ mg^-1^ protein. The data shown are the average of triplicate determination ± standard deviation

|  | DON | CUL |
| --- | --- | --- |
| UGT | Specific activity (nmol min<sup>-1</sup> mg<sup>-1</sup>) |  |
| BdUGT5g03300 | 1.07 ± 0.03 | $\approx 1 \times 10^{-3}$ |
| HvUGT13248 | 8.77 ± 0.52 | 6.94 ± 0.52 |
| OsUGT79 | 159 ± 33 | 1.85 ± 0.05 |
| OsUGT79 Q202A* | 161 ± 46 | 2.23 ± 0.08 |
| OsUGT79 H122A/L123A/Q202A* | 6.60 ± 0.31 | 2.89 ± 0.10 |

\* Active site mutants of OsUGT79 (Wetterhorn *et al.*, 2017)

OsUGT79 glucosylated CUL at a low rate of 1-2% compared to DON (**Table 2**). We also investigated two active site mutants that were previously created to expand the substrate range of OsUGT79 with trichothecenes. With DON, OsUGT79 Q202A was reported to behave kinetically similar to the wildtype enzyme, while the triple mutant OsUGT79 H122A/L123A/Q202A had reduced affinity for DON, but also the broadest substrate range and accepted C4-acetylated trichothecenes (Wetterhorn *et al*., 2017). Herein, wild type OsUGT79 and both mutants displayed similar conversion rates with CUL, while the catalytic rates observed with DON clearly differed (**Table 2**).

### Inhibition kinetics of UGTs with DON and CUL

To gain further insights into the mechanism of UGT inhibition by CUL, we tested the influence of CUL on the reaction kinetics of all UGTs with DON and UDP-glucose. It should be noted that the used assay quantifies released UDP *via* an enzymatic cascade causing NADH oxidation) and does not distinguish between glucosylation of DON or CUL. To compensate for that, the blank reactions (“0 mM DON”) also contained CUL. The such estimated values for the inhibition constants of competitive (*K*_ic_, equation 3) and uncompetitive inhibition (*K*_iu_, equation 4) are displayed in **Table 3**.

**Table 3.** Influence of culmorin on the kinetics of UDP-glucosyltransferases with deoxynivalenol and UDP-glucose. Kinetic parameters and constants of competitive (*K*_ic_) and uncompetitive (*K*_iu_) inhibition are shown. Enzyme assays were performed at pH 7, 25 °C using a coupled enzyme assay to detect released UDP via NADH oxidation.

|  | Deoxynivalenol (DON)<sup>a</sup> |  |  |  | UDP-glucose<sup>b</sup> |  |
| --- | --- | --- | --- | --- | --- | --- |
| | $V_{max}$<br>($\mu\text{mol min}^{-1} \text{mg}^{-1}$) | $K_M$<br>(mM DON) | $K_{ic}$<br>($\mu\text{M CUL}$) | $K_{iu}$<br>($\mu\text{M CUL}$) | $K_M$<br>(mM UDP-gluc) | $K_{iu}$<br>($\mu\text{M CUL}$) |
| BdUGT5g03300 | 0.14 | 1.1 | < 1 | - | 0.19 | 1.4 |
| HvUGT13248 | 0.18 | 1.8 | 56 | - | 0.33 | 91 |
| OsUGT79 | 0.63 | 0.19 | 146 | 208 | 0.21 | 124 |
| OsUGT79 Q202A <sup>c</sup> | 0.85 | 0.29 | 16 | 56 |  |  |
| OsUGT79 H122A/L123A/Q202A <sup>c</sup> | 0.26 | 1.3 | 65 | 217 |  |  |

<sup>a</sup> 1 mM UDP-glucose, 1-17 mM DON
<sup>b</sup> 5 mM DON, 1-5 mM UDP-glucose
<sup>c</sup> Active site mutants of OsUGT79 (Wetterhorn *et al.*, 2017)

BdUGT5g03300 activity with DON was competitively inhibited with an about 50-fold increase of apparent *K*_M_ with DON at already 10 µM CUL (**Fig. 3A** and **3B**). *K*_ic_ was estimated in the low µM range (**Table 3**). Compared to the *K*_M_ of 1.1 mM DON, this implies remarkably strong binding of CUL. Inhibition kinetics with UDP-glucose as the variable substrate (5 mM DON) indicated an uncompetitive mechanism (**Fig. 3C**) with a decrease of both *V*_max_ and *K*_M_. *K*_iu_ with UDP-glucose was estimated at 1.4 µM CUL and lies in the same concentration range as *K*_ic_ obtained with DON. The corresponding model of uncompetitive inhibition (equation 4) assumes that the uncompetitive inhibitor binds only to the enzyme-substrate complex (UDP-glucose bound to BdUGT5g03300). Accordingly, our results imply that CUL binds with high affinity (*K*_D_ ≈ 1 µM range) in an unproductive position/orientation at the acceptor binding site of BdUGT5g03300 when UDP-glucose is already present at the donor-binding site.

**Fig. 3.**
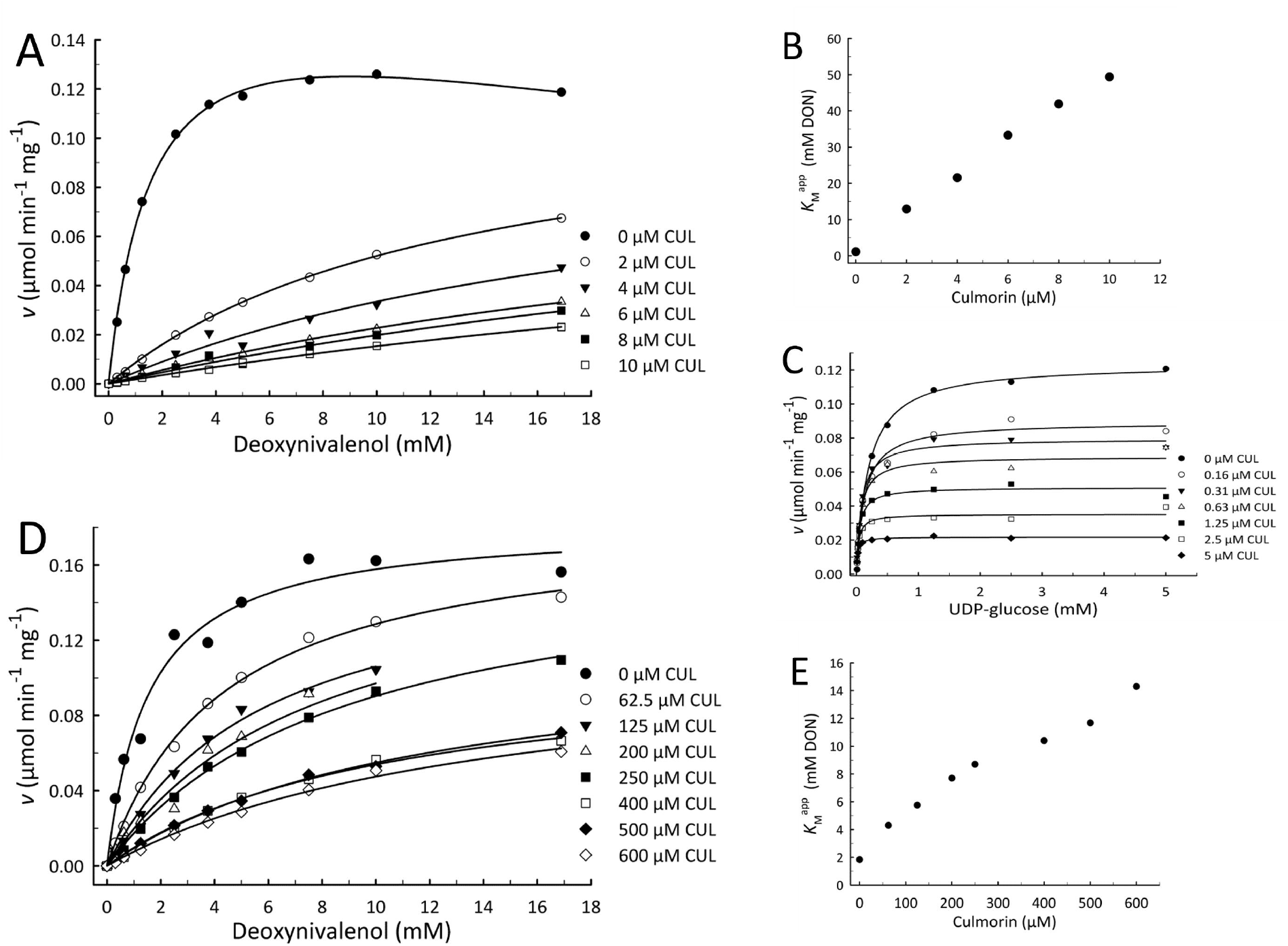
Influence of culmorin (CUL) on the kinetics of deoxynivalenol (DON) glucosylation by the UDP-glucosyltransferases BdUGT5g03300 and HvUGT13248. **(A)** BdUGT5g03300, inhibition kinetics of CUL with 1 mM UDP-glucose, DON concentration variable. **(B)** BdUGT5g03300, apparent *K*_M_ values with DON plotted against CUL concentration. **(C)** BdUGT5g03300, inhibition kinetics of CUL with 5 mM DON, UDP-glucose variable. **(D)** HvUGT13248, inhibition kinetics of CUL with 1 mM UDP-glucose, DON concentration variable. (E) HvUGT13248, apparent *K*_M_ with DON plotted against CUL concentration.

Consistent with the above observation that CUL is conjugated by HvUGT13248 with comparable efficiency as with DON, inhibition of DON-glucosylation clearly followed a competitive mechanism with an about 10-fold increase of apparent *K*_M_ of DON at 600 µM CUL (**Fig. 3D** and **3E**). The estimated *K*_ic_ (56 µM CUL) lies in a similar concentration range as *K*_M_ with CUL (25 µM). As observed with BdUGT5g03300, inhibition was uncompetitive with UDP-glucose (**Table 3**).

OsUGT79 has low activity with CUL and is only moderately inhibited by CUL. The case is kinetically interesting as in assays without CUL, OsUGT79 displays pronounced substrate inhibition by DON (**Fig. 4A**). The Haldane model (uncompetitive substrate inhibition, equation 2) is usually used to fit such data. It is assumed that the substrate can bind to the already formed enzyme-substrate complex at a second site and acts as inhibitor, for example by preventing the product from leaving (Bisswanger, 2015). Interestingly, the observed substrate inhibition by DON was reduced with increasing CUL concentrations (**Fig. 4A**), suggesting that CUL competes with DON for the hypothetical substrate inhibition site. CUL further caused a clear decrease in predicted *V*_max_. Although a slight increase of *K*_M_ was indicated (**Fig. 4B**), this remained within a narrow concentration range and *K*_M_ remained effectively unchanged. This behavior resembles non-competitive inhibition, a case of mixed inhibition (equation 5) where the inhibitor binds independently of whether the substrate is bound at the active site (*K*_ic_ ≈ *K*_iu_). The *K*_i_ values of CUL obtained with OsUGT79 (**Table 3**) would confirm this model, although a completely non-competitive mechanism is unlikely, as CUL is also a substrate of OsUGT79. As in the other two cases (HvUGT13248, BdUGT5g03300), inhibition with respect to UDP-glucose followed an uncompetitive mechanism. Compared to wild type OsUGT79, even stronger binding of CUL by OsUGT79 Q202A was indicated by the estimated *K*_i_ values and the competitive component is clearly increased in OsUGT79 Q202A (Table 3). The triple mutant OsUGT79 H122A/L123A/Q202A did not show substrate inhibition by DON (*K*_M_ = 1.3 mM DON) in the investigated concentration range (≤ 16 mM DON) and here, the competitive character of inhibition by CUL clearly dominates (**Table 3**).

**Fig. 4.**
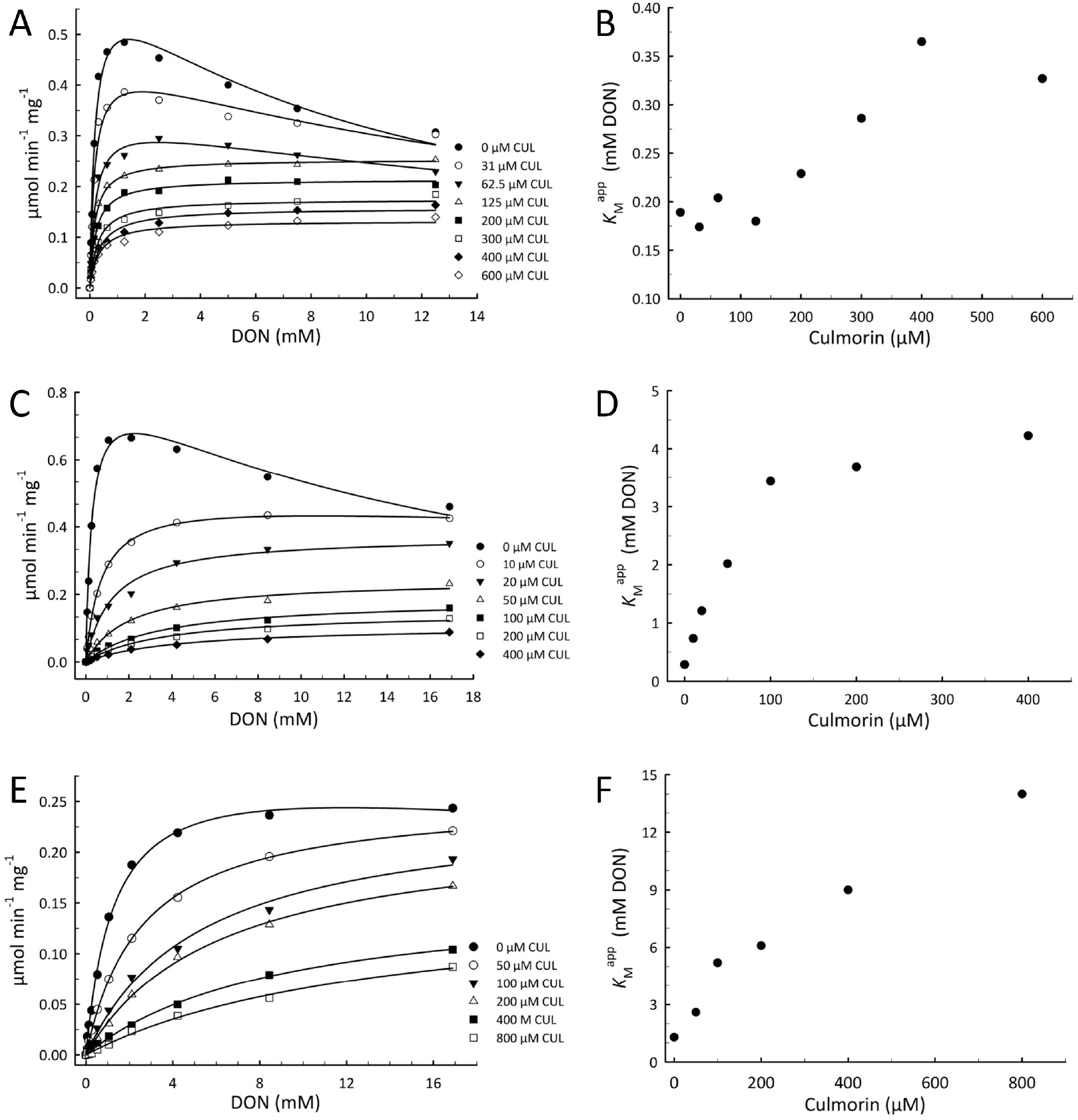
Influence of culmorin (CUL) on the kinetics of deoxynivalenol (DON) glucosylation by OsGT79. **(A)** OsUGT79 wild type, kinetic plots with 1 mM UDP-glucose, deoxynivalenol (DON) variable. **(B)** apparent *K*_M_ of DON plotted against CUL concentration. **(C)** OsUGT79 Q202A, kinetic plots with 1 mM UDP-glucose, DON variable. **(D)** apparent *K*_M_ of DON plotted against CUL concentration. **(E)** OsUGT79 H122A/L123A/Q202A, kinetic plots with 1 mM UDP-glucose, DON variable. **(F)** apparent *K*_M_ of DON plotted against CUL concentration.

### Is inhibition of BdUGT5g03300 by CUL responsible for the synergy of CUL with DON in *B. distachyon?*

*Because o*f the strong inhibition of BdUGT5g03300 by CUL and the fact that BdUGT5g03300 is crucial for Fusarium/DON resistance in *B. distachyon* (Pasquet *et al*., 2016), we further investigated the combined effect of DON and CUL on *B. distachyon*. This should clarify whether the effect observed on BdUGT5g03300 *in vitro* can be connected to DON-glucosylation *in vivo*. Compared to Apogee wheat, *B. distachyon* Bd21-3 appears more tolerant to DON. While the former already showed impaired root growth at 3 mg L^-1^ DON (**Fig. 2**), Bd21-3 was not or barely affected between 2-4 mg L^-1^ DON. No more root growth was observed at 20 mg L^-1^ DON (**Supplementary Fig. S3**). As observed with wheat, the presence of only CUL did not elicit any significant effect on root length. However, CUL caused a clear dose-dependent synergistic effect with DON when applied at concentrations up to 2 mg L^-1^ at 2 mg L^-1^ DON (**Fig. S3, Supplementary Table S6**).

To investigate the potential involvement of BdUGT5g03300, we analyzed DON and CUL glucosylation in the roots of *B. distachyon* TILLING mutant 6829-7 carrying a truncated (W345^*^) BdUGT5g03300 gene (Pasquet *et al*., 2016). As control line, the TILLING line 6829-3 carrying the wild type BdUGT5g03300 allele was used. Roots were incubated in MS medium containing DON (50 mg L^-1^) and/or CUL (10 mg L^-1^) for 24 h. **Fig. 5 A** shows that in line 6829-7 (BdUGT5g03300 W345^*^), D3G synthesis was low and not further reduced in presence of CUL. In TILLING line 6829-3 (wild type BdUGT5g03300 allele), D3G synthesis was clearly functional but suppressed in the presence of CUL, with D3G concentrations at DON+CUL treatment similar to those in the W345^*^ mutant (**Fig. 5B**). This indicates that BdUGT5g03300 is mainly responsible for D3G synthesis and suggests its inhibition in the presence of CUL *in vivo*. We further investigated the BdUGT5g03300 over-expressing line OE-10R14 that was previously shown to be highly DON/Fusarium resistant (Pasquet *et al*., 2016). In root elongation assays, this line tolerated up to 50 mg L^-1^ DON without significant reduction in root length. Remarkably, addition of CUL at even low levels (0.5 mg L^-1^) to 50 mg L^-1^ DON caused a severe breakdown of resistance to DON (**Fig. 5C, Supplementary Table S7**). Analysis of root extracts (6 and 24 h incubation) indicated high D3G formation capacity in the overexpression line (**Fig. 5D**). The presence of CUL reduced D3G synthesis, but not as drastically as would have been expected (about 58.6% at 24 h, p = 0.042). However, assays were analyzed at different time points as the root elongation assays (6 days). More detailed investigations may clarify the time-dependent kinetics of DON/CUL glucosylation in the overexpression line. Nevertheless, these results clearly suggest that the tolerance gained by overexpression of BdUGT5g03300 is reduced in the presence of CUL.

**Fig.5.**
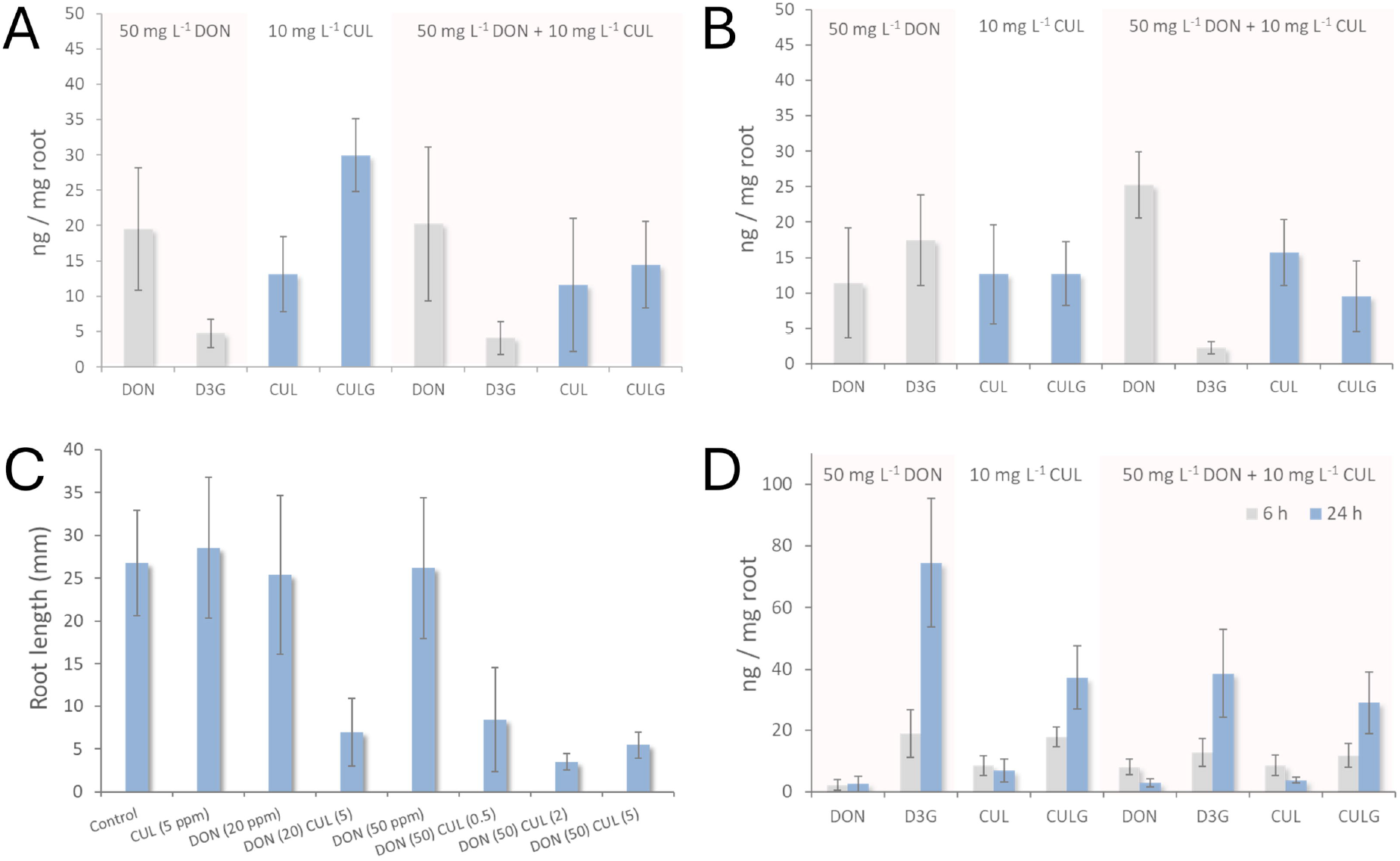
Impact of deoxynivalenol (DON) and culmorin (CUL) on roots of *B. distachyon* seedlings. (**ABD**) DON, DON-3-glucoside (D3G), CUL and CUL-glucoside (CULG) concentrations in *B. distachyon* roots extracted after incubation in liquid Murashige-Skoog (MS) medium with 50 mg L^-1^ (169 µM) DON and/or 50 mg L^-1^ (210 µM) CUL. **(A)** Analysis of root extracts of TILLING mutant 6829-7 (truncated BdUGT5g03300 gene, W345^*^), after 24 h incubation with DON and/or CUL (n = 8). **(B)** Analysis of root extracts of TILLING mutant 6829-3 carrying wild-type BdUGT5g03300 allele incubated for 24 h with DON and/or CUL (n = 8). **(C)** average total root length (n ≈ 10, error bars indicate SD) of BdUGT5g03300 over-expression line OE10R14 seedlings after 6 days of growth in MS agarose medium containing different concentrations of DON and CUL; concentrations in mg L^-1^ are given in parentheses. **(D)** Analysis of root extracts of BdUGT5g03300 over-expression line OE10R14 after incubation for 6 and 24 h in liquid MS with DON and/or CUL (n=5).

### BdUGT5g03370 and BdUGT5g03380 glucosylate CUL

The above results (**Fig. 5**) have further indicated that *Brachypodium* is able to glucosylate CUL and suggest that BdUGT5g03300 is not responsible. TILLING line 6829-7 (BdUGT5g03300 W345^*^) showed even higher CUL glucosylation than line 6829-3 carrying the intact BdUGT5g03300 allele (**Fig 5B**). BdUGT5g03300 is member of a DON/Fusarium responsive gene cluster containing six UGT genes designated BdUGT5g02780, BdUGT5g03300, BdUGT5g03370, BdUGT5g03380, BdUGTg03390 and BdUGT5g03400 (Schweiger *et al*., 2013a). Of these, only BdUGT5g03300 possesses activity with DON. BdUGT02780 is active with NIV but not with DON, the functions of the other cluster members are unknown (Schweiger *et al*., 2013a). Tentative assays with protein extracts of yeast expressing these UGTs (Schweiger *et al*., 2013a) indicated that BdUGT5g03370 and BdUGTg03380, both highly expressed upon DON treatment/Fusarium infection (Schweiger *et al*., 2013a) are capable of glucosylating CUL (data not shown). We have further expressed both enzymes in *E. coli* and estimated the kinetics of the one-step purified enzymes, showing that BdUGT5g03370 glucosylates CUL with a *K*_M_ of 59 ± 7 µM CUL and *V*_max_ of 0.57 ± nmol min^-1^ mg^-1^; BdUGT5g03380 with *K*_M_ 150 µM CUL and V_max_ 0.43 ± 0.03 nmol min^-1^ mg^-1^ (**Supplementary Fig. S4**).

## Discussion

Culmorin is a somewhat enigmatic *Fusarium* metabolite as it persistently co-occurs with DON at high levels but does not appear to possess an obvious function. The fact that expression of the relevant biosynthetic genes of both metabolites is co-regulated under trichothecene inducing conditions strongly suggests a functional link. Furthermore, CUL originates from the same biosynthetic pool as DON, and it is reasonable to hypothesize a complementary function. Previous studies already indicated a possible synergy of CUL with DON in plants and hypothesized that CUL may interfere with DON-detoxification (Dowd *et al*., 1989; Wang and Miller, 1988; Wipfler *et al*., 2019). A previously published collaborative study reported that CUL suppressed DON-glucuronidation by human glucuronosyltransferases in liver microsomes (Woelflingseder *et al*., 2019). Here, we provide evidence that CUL-deficient *F. graminearum* strains are less aggressive on wheat. We further show that the presence of CUL interferes with the detoxification of DON by conjugation to D3G in both wheat and *B. distachyon*, indicating that CUL may contribute to *Fusarium* virulence by acting synergistically with DON.

Glucosylation is an important phase II conjugation reaction in plant secondary metabolism that alters solubility and chemical activity of bioactive compounds. It is involved in homeostasis of endogenous metabolites including defense metabolites and phytohormones, but also in the detoxification of xenobiotics (Bowles *et al*., 2006; Gachon *et al*., 2005; Lim and Bowles, 2004). Plant glucosylation of small molecules is catalyzed by UGTs assigned to family 1 in the carbohydrate active enzyme database CAZy (http://www.cazy.org), which currently lists 138 glycosyltransferase families (Campbell *et al*., 1997; Coutinho *et al*., 2003). Family 1 UGTs are composed of two Rossman folds and the active site is located in a cleft between the two domains (Lairson *et al*., 2008; Qasba *et al*., 2005). UGTs are usually highly specific for the donor substrate (“glycon”; i.e., UDP-glucose) that binds to the highly conserved plant secondary product glycosyltransferase (PSPG) motif located at the *C*-terminal domain, considered the signature motif of family 1 UGTs (pfam00201). The less conserved *N*-terminal domain interacts with the acceptor substrate (aglycon). Although some members are specific for the acceptor substrate, UGTs are generally promiscuous towards the aglycon (Gachon *et al*., 2005; Osmani *et al*., 2009), a prerequisite for catalysis is the correct positioning of the functional group amenable for glycosylation (Osmani *et al*., 2009).

DON-3-glucoside (D3G), previously considered a “masked mycotoxin”, frequently co-occurs with its parental toxin to a variable extent (Berthiller *et al*., 2013), indicating that glucosylation of DON is a relevant plant defense mechanism. Several plant UGTs with the ability to convert DON into the non-toxic D3G have been identified (Gharabli *et al*., 2023; Michlmayr *et al*., 2018). Following our observation that CUL inhibits DON-glucosylation in wheat, a future research question will be to investigate wheat UGTs regarding their inhibition by CUL. However, although several candidate UGTs of hexaploid wheat (*Triticum aestivum*) have been proposed to be involved in Fusarium resistance, there is little evidence for their actual contribution to DON detoxification. The DON-responsive TaUGT3 was reported to increase tolerance to DON (Lulin *et al*., 2010), but DON-glucosylation could not be confirmed with a DON-sensitive yeast strain (Schweiger *et al*., 2010). Another pathogen inducible UGT, TaUGT12887, was found only weakly active with DON (Schweiger *et al*., 2013b). Traes_2BS_14CA35D5D, a homolog of BdUGT5g03300 provides Fusarium resistance in planta, but has not yet been biochemically characterized with DON (Gatti *et al*., 2018b). TaUGT6 is another candidate proposedly involved in resistance but the purified protein was described as only “active to some extent” with DON (He *et al*., 2020).

In the absence of biochemically validated DON-detoxifying UGTs from wheat, we herein investigated the previously characterized BdUGT5g03300 (B. distachyon), OsUGT79 (rice) and HvUGT13248 (barley) as these possess sufficient activities with DON to conduct kinetic assays (Michlmayr *et al*., 2018). Barley HvUGT13248 is DON-inducible (Gardiner *et al*., 2010) and confers resistance when over-expressed in *Arabidopsis thaliana* and wheat (Li *et al*., 2017; Li *et al*., 2015; Mandalà *et al*., 2019). Loss of function of HvUGT13248 leads to reduced Fusarium spreading resistance in barley, demonstrating the important role of toxin detoxification in host defense (Bethke *et al*., 2023). Also, BdUGT5g03300 (“Bradi5g03300”) is important for Fusarium resistance of the monocot model plant *B. distachyon* (Pasquet *et al*., 2016) and provided resistance to wheat upon heterologous over-expression (Gatti *et al*., 2018a). Disruption of BdUGT5g03300 in *B. distachyon* by TILLING led to increased susceptibility to DON and reduced resistance against *F. graminearum* (Pasquet *et al*., 2016). OsUGT79 from rice was identified by homology to both BdUGT5g03300 and HvUGT13248. It is highly active with DON and its 3D structure yielded crucial insights into trichothecene accommodation at the active site (Wetterhorn *et al*., 2017; Wetterhorn *et al*., 2016). We therefore used these UGTs as case-study models to investigate whether CUL is capable of impeding DON-detoxification by serving as alternative substrate or inhibitor. This indicated that CUL can in fact interfere with DON-detoxification by UGTs, although the effect is clearly variable in magnitude and mechanism. Inhibition by CUL was uncompetitive with UDP-glucose in all cases which indicates that the inhibitor can only bind when the enzymes are already present in complex with UDP-glucose. This is consistent with the fact that UGTs undergo conformational changes upon substrate binding and the sugar donor usually binds first (Breton *et al*., 2012). It is possible that UDP-glucose induces conformational changes to make the corresponding sites accessible.

Our results further suggest that CUL inhibits UGTs by binding to sites that recognize DON although it does not necessarily mimic DON as substrate analogue. This results in remarkably strong unproductive binding/inhibition of BdUGT5g03300 and competitive inhibition of HvUGT13248 by CUL serving as alternative substrate. In the case of OsUGT79, CUL appears to compete with the alternative binding site that DON occupies to exert substrate inhibition. Judged from the logP values published on PubChem (https://pubchem.ncbi.nlm.nih.gov/, accessed Sept, 2025), CUL is strongly hydrophobic (XLogP3-AA = 3.3) while DON (XLogP3-AA = -0.7) is hydrophilic. Considering the functional “promiscuity” of defense related UGTs and the largely hydrophobic nature of UGT acceptor binding pockets (Osmani *et al*., 2009), it may not be surprising that a hydrophobic metabolite of relatively small mass (CUL 238 g/mol) can bind, in some cases even with high affinity, to the active sites of a UGT in a productive or unproductive orientation. From such a perspective, it can be hypothesized that other phase II detoxification enzymes such as glutathione transferases with low selectivity as “functional principle” could be affected analogously. We have previously shown that tau class GSTs from wheat are capable of conjugating DON as well (Michlmayr *et al*., 2025). Besides DON detoxification, both UGTs and GSTs play diverse catalytic roles in plant stress responses and may be likewise be affected by CUL. As demonstrated here, both wheat and *B. distachyon* are able to glucosylate CUL, which would for example diminish its inhibitory activity on HvUGT13248. In contrast, also the CUL-glucosides were inhibitory to BdUGT5g03300, which implies that glucosylation catalyzed by other UGTs such as BdUGT5g03370 and BdUGT5g03380 may not counteract its synergistic effect with DON. With respect to transgenic approaches to increase Fusarium resistance by UGT overexpression, this implies that OsUGT79 or HvUGT13248 may have advantages over BdUGT5g03300.

In conclusion, this study presents biochemical evidence that CUL may contribute to *Fusarium* virulence by inhibiting DON-detoxification by plant UGTs, resulting in a synergistic effect with DON. CUL production could allow the fungus to overcome transgenic or breeding efforts to increase Fusarium resistance by increasing DON-detoxification by glucosylation. The warfare between trichothecenes as *Fusarium* virulence factors and plant detoxification enzymes is presumably ongoing since about 27 million years (O’Donnell *et al*., 2013). Our results imply that UGTs of different plant species not only had to co-evolve with structurally diverse virulence factors but also with co-produced inhibitors such as culmorin, and potentially many other compounds.

## Abbreviations

(CUL): culmorin
(DON): deoxynivalenol
(D3G): deoxynivalenol-3-glucoside
(TILLING): targeting induced local lesion in genomes
(UGT): UDP-glucosyltransferase

## Supplementary data

**Table S1**. Production of DON and CUL by *F. graminearum* PH-1 and two *CLM1 knock-out t*ransformants on 2-SM medium

**Table S2**. Statistical analysis of wheat infection scores

**Table S3**. DON and D3G concentrations in infected wheat heads

**Table S4**. Statistical analysis of primary wheat root length after DON and CUL treatment.

**Table S5**. Statistical analysis of total wheat root length after DON and CUL treatment.

**Table S6**. Statistical analysis of *B. distachyon* (Bd21-3) total root length after DON and CUL treatment.

**Table S7**. Statistical analysis of total root length of BdUGT5g03300 over-expression *B. distachyon* line OE10R14 after DON and CUL treatment.

**Fig. S1**. Experimental set-up of root elongation assays with Apogee seedlings.

**Fig. S2**. Kinetics of HvUGT13248 with culmorin.

**Fig. S3**. *B. distachyon* (Bd21-3) seedlings after exposure to DON and CUL.

**Fig. S4**. Kinetics of BdUGT5g03370 and BdUGT5g03380 with CUL.

## Acknowledgements

We thank Prof. Marc Lemmens for providing DON and Prof. Ivan Rayment for providing the OsUGT79 mutants. We also thank Ass. Prof. Elisabeth Varga for analytical support.

## CRediT Authorship statement

HM, GW, FB & GA: Conceptualization; HM: Formal Analysis; HM, GA, RK & FB: Funding Acquisition; HM, GW, KR, MS, KF, AM, MD & FB: Investigation; KR & MS: Methodology; PF, JW, & MD: Resources; GW, RK, FB & GA: Supervision; HM & GW: Writing – Original Draft Preparation; All Authors: Writing – Review & Editing

## Data availability

The data supporting the conclusions of this article are included within the article and its supplementary data published online.

